## Supplemental Figures S1-5 for "Genome sequence and characterization of five bacteriophages infecting *Streptomyces coelicolor* and *Streptomyces venezuelae*: Alderaan, Coruscant, Dagobah, Endor1 and Endor2"

**Supplementary data**

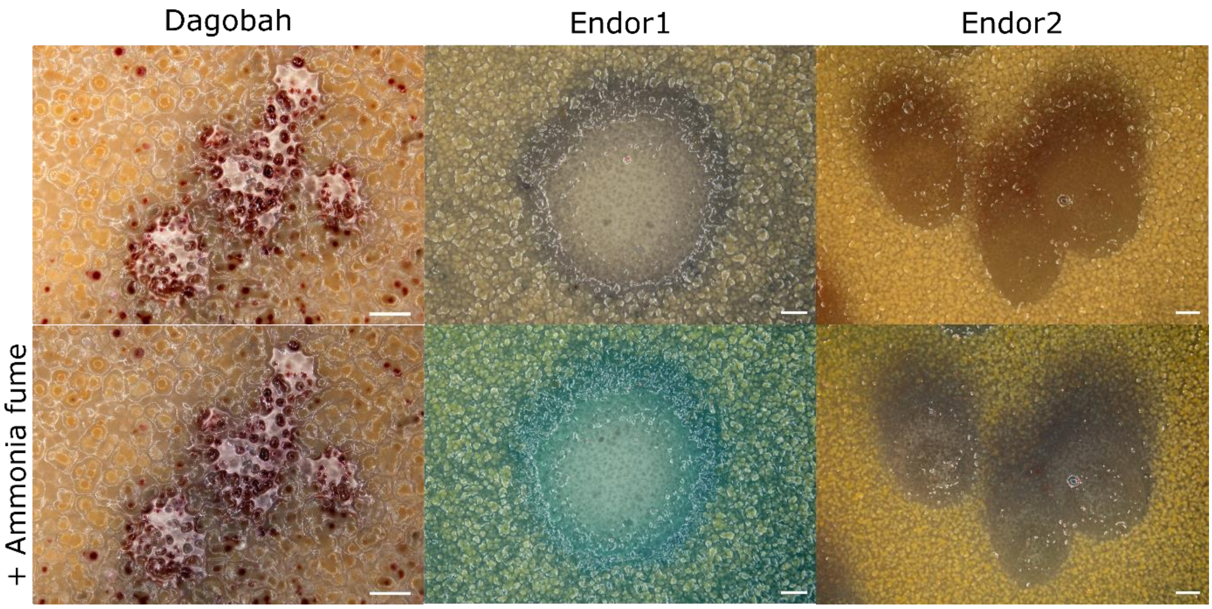


**Figure S1**. Close-ups of phage plaques imaged using a Nikon SMZ18 stereomicroscope, before (upper row) and after (lower row) exposure to ammonia fumes. *S. coelicolor* M145 was infected by phages using GYM double agar overlays. The plates were incubated at 30°C overnight and then kept at room temperature for two (Dagobah and Endor2) or three days (Endor1). The ammonia fume test was performed as follows: the plates were inverted and exposed to ammonia fumes for 15 min by placing 5 ml of 20% ammonium hydroxide solution on the inner surface of the lid. Scale bar: 1 mm.

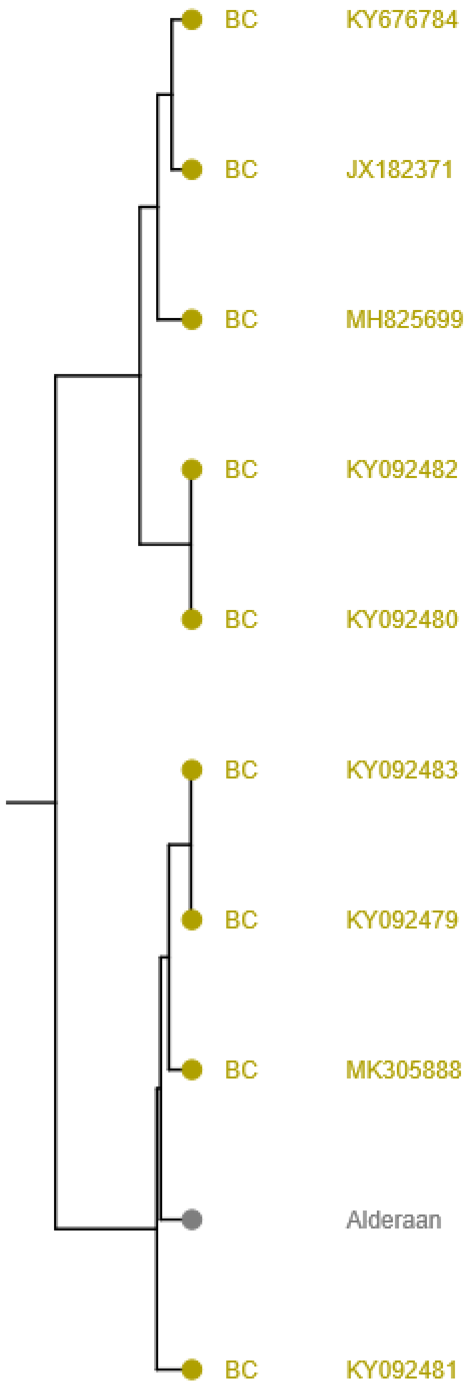


**Figure S2**. Subclade dendrogram with *Streptomyces* phage Alderaan and its closely related actinophages (enlargement from Figure 5).

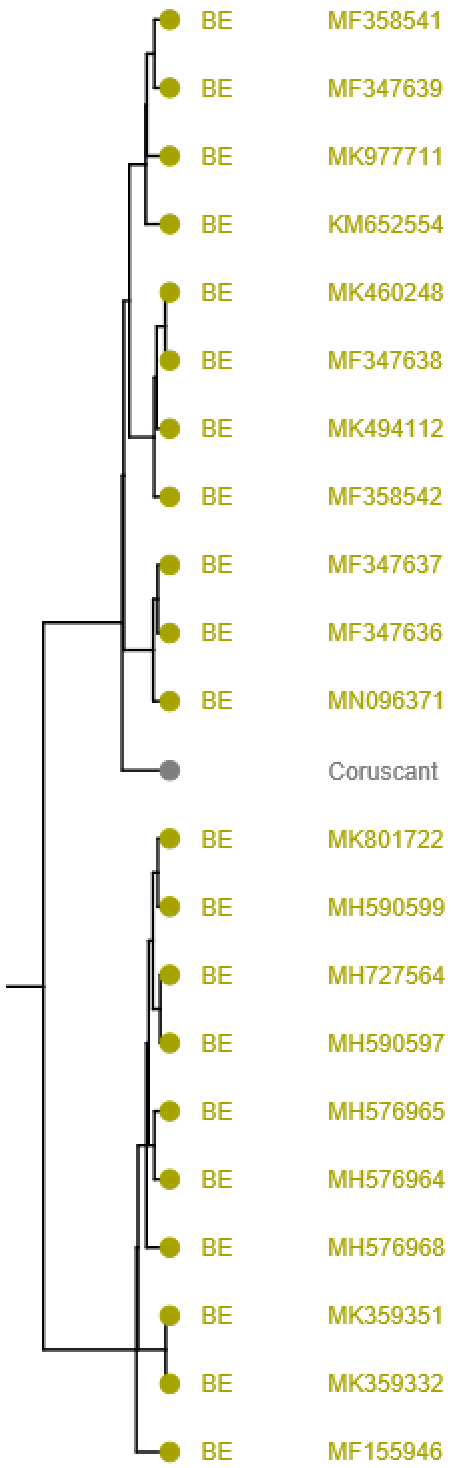


**Figure S3.** Subclade dendrogram with *Streptomyces* phage Coruscant and its closely related actinophages (enlargement from Figure 5).

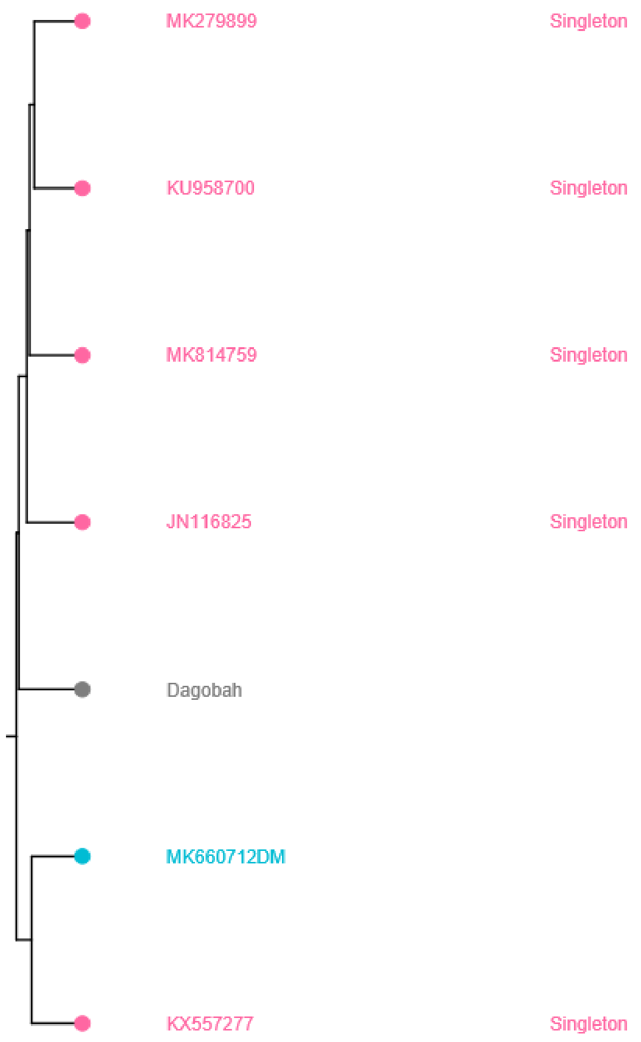


**Figure S4.** Subclade dendrogram with *Streptomyces* phage Dagobah and its closely related actinophages (enlargement from Figure 5).

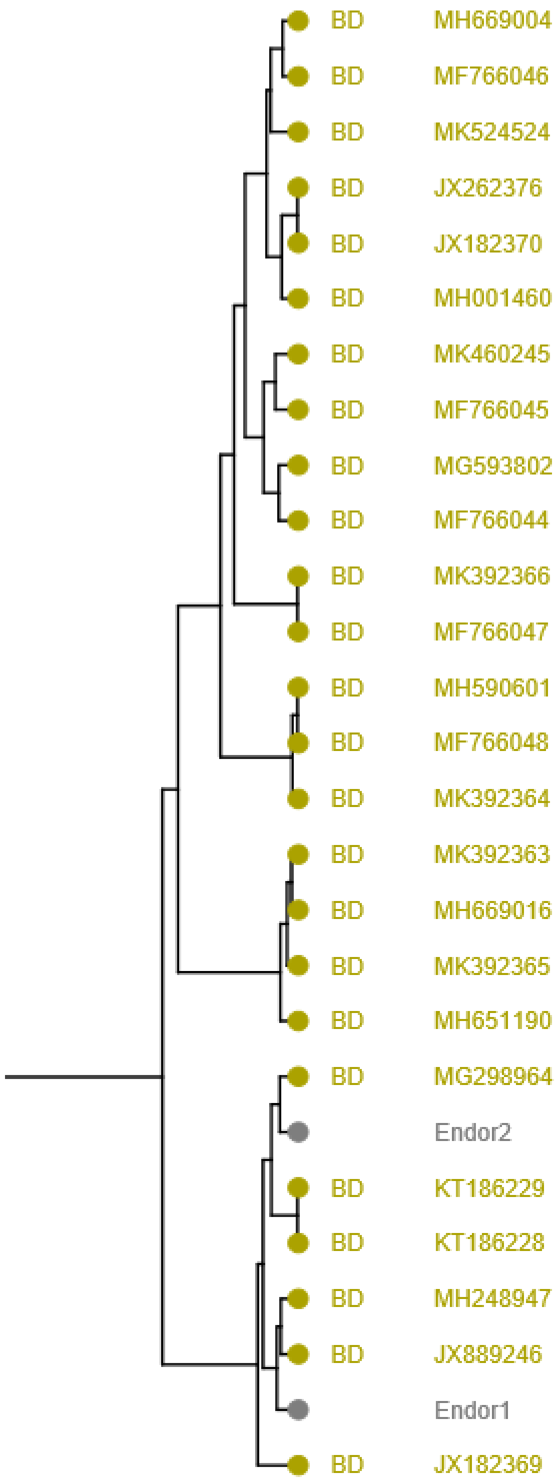


**Figure S5.** Subclade dendrogram with *Streptomyces* phages Endor1 and Endor2 and their closely related actinophages (enlargement from Figure 5).
